# NMDA receptor misalignment in iPSC-derived neurons from a multi-generational family with inherited Creutzfeldt-Jakob disease

**DOI:** 10.1101/2022.05.20.491674

**Authors:** Nhat T.T. Le, Robert C.C. Mercer, Aldana D. Gojanovich, Alice Anane, Seonmi Park, Bei Wu, Pushpinder S. Bawa, Regeneron Genetics Center, Gustavo Mostoslavsky, David A. Harris

## Abstract

The most common subtype of genetic prion disease is caused by the E200K mutation of the prion protein. We have obtained samples from 22 members of a multi-generational Israeli family harboring this mutation, and generated a library of induced pluripotent stem cells (iPSCs) representing nine carriers and four non-carriers. Whole-exome sequencing was performed on all individuals. A comparison of neurons derived from E200K iPSCs to those from non-carriers revealed the presence of several disease-relevant phenotypes. Neurons from E200K carriers were found to contain thioflavin S-positive accumulations of PrP in their cell bodies. In addition, these neurons displayed disruptions of NMDA receptor/PSD95 co-localization at postsynaptic sites. Our study shows that iPSC-derived neurons, which express physiologically relevant levels of mutant PrP in a human neuronal context, can model certain aspects of human prion disease, offering a powerful platform for investigating pathological mechanisms and testing potential therapeutics.

## Introduction

Prion diseases are invariably fatal neurodegenerative conditions affecting a wide range of mammalian species. This group includes diseases such as bovine spongiform encephalopathy (BSE or “mad cow disease”), scrapie, which afflicts sheep and goats, and chronic wasting disease (CWD) which affects cervid populations in North America, Asia and Europe. In humans, prion diseases include Creutzfeldt-Jakob disease (CJD), Gerstmann-Sträussler-Scheinker disease (GSS), fatal familial insomnia (FFI), and kuru (Mercer et al., 2018; Prusiner, 1998). The key molecular event underlying these disorders is a structural transformation of the cellular prion protein (PrP^C^), a normal cell-surface glycoprotein, into a pathogenic, amyloid conformation, denoted PrP^Sc^ (Kraus et al., 2021; Manka et al., 2022). This structural rearrangement endows PrP^Sc^ with the ability to catalyze its own formation through a self-templating mechanism, and imparts upon it distinct biochemical attributes, including resistance to protease digestion and insolubility in detergents (Meisl et al., 2021; Mercer *et al*., 2018). In humans, the etiology of these diseases is threefold; i) sporadic CJD is by far the most common human prion disease and accounts for ~85% of all cases, ii) a small proportion of cases are acquired by infection through iatrogenic means or through consumption of contaminated tissues, and iii) ~15% of cases are inherited in an autosomal dominant manner due to point and insertion/deletion mutations in *PRNP*, the gene encoding the prion protein (Wadsworth and Collinge, 2007).

The most prevalent prion disease-associated mutation in humans is a substitution of glutamic acid for lysine at residue 200 (E200K), which causes CJD. This is a highly pathogenic mutation, displaying almost 100% penetrance (Minikel et al., 2016). CJD-E200K patients most commonly present with dementia, ataxia and myoclonus; other symptoms such as pyramidal signs, Parkinsonism and dystonia are less frequently present. Spongiform change, neuronal loss, and gliosis predominate in the neocortex, basal ganglia, thalamus, and substantia nigra. Diffuse synaptic immunostaining of PrP is present in virtually all CJD-E200K patients. However a more plaque-like deposition predominates in individuals who are heterozygous (methionine/valine) at the polymorphic residue 129 (Kovacs et al., 2011). The identity of this residue is known to have profound effects upon the presentation and/or disease course of many human prion diseases, however neither the clinical signs nor pattern of spongiosis associated with CJD-E200K appear to be influenced by this polymorphism (Goldfarb et al., 1992; Kovacs *et al*., 2011). Pathological tau deposition can also be found in more than 90% of CJD-E200K patients (Kovacs *et al*., 2011).

Several experimental systems have been used previously to model inherited prion diseases, including those caused by the E200K mutation. Work from our laboratory demonstrated that, when overexpressed in CHO cells, E199K PrP (the mouse homologue of the E200K human mutation) acquired some of the biochemical properties of prions, including detergent insolubility and resistance to protease digestion (Lehmann and Harris, 1996a; b). However, immortalized cells (perhaps with one exception (Schätzl et al., 1997)) do not exhibit signs of toxicity upon prion infection, making them unsuitable to examine the mechanisms underlying prion-induced cytopathology (Mercer and Harris, 2019; 2022). Extensive research has focused on the development of E200K transgenic mice, but none of the existing models fully recapitulate all features of the disease. Mice expressing the mutated human protein do not develop disease (Asante et al., 2009), and while mice expressing the mutation in the context of a mouse/human hybrid PrP do succumb to a neurodegenerative syndrome, pathological changes such as vaccuolation and PrP^Sc^ deposition are minor and protease resistant PrP is not typical of that found in CJD-E200K (Friedman-Levi et al., 2011; Friedman-Levi et al., 2013). Human iPSCs expressing wild-type PrP have been used to generate cerebral organoids (COs), which were shown to be capable of propagating sporadic CJD prions (Groveman et al., 2019). While COs generated from iPSCs expressing E200K PrP did not contain any detectable PrP^Sc^, and displayed no PrP or tau-related pathology (Foliaki et al., 2020; Gonzalez et al., 2018; Smith et al., 2022), neuronal network communication has been reported to be disrupted (Foliaki et al., 2021). Hyperphosphorylated tau has been noted in iPSC-derived neurons expressing PrP with a substitution of tyrosine for asparagine at residue 218 (Y218N), which is associated with GSS, demonstrating that at least some aspects of inherited prion disease can be modeled using iPSC derived systems (Matamoros-Angles et al., 2018).

Here, we describe a library of iPSCs obtained from an Israeli family harboring the E200K PrP mutation. To our knowledge, this is the first report of iPSC lines derived from multiple members of a single, multi-generational family carrying a prion disease-related mutation. We have analyzed cortical pyramidal neurons derived from these iPSCs, and find that neurons from E200K carriers display specific biochemical and synaptic abnormalities not observed in neurons from non-carriers. We also performed whole-exome sequencing of genomic DNA from each member of this cohort. This new resource represents a powerful experimental platform for dissection of the molecular mechanisms underlying human prion diseases.

## Results

### Molecular analysis of donor PRNP

The Israeli family, of Libyan Jewish origin, spans four generations. We consented 22 individuals in generations 2-4, including 11 carriers of the E200K PrP mutation, as well as 11 non-carriers (Table 1; ages, genders, and lineage are not shown to preserve anonymity). None of the living carriers have shown symptoms of CJD, but one carrier (CJD40) who died at the age of 99 had displayed mild dementia for two years before death and subsequent to donation. Sanger sequencing of the *PRNP* gene was performed to determine carrier status of the E200K mutation and haplotype phasing with other *PRNP* polymorphisms. Among non-carriers, the majority (9/11) are heterozygous for the common M/V polymorphism at codon 129. The other two individuals are M/M and V/V homozygotes. The E200K mutation was heterozygous in all carriers, and was found exclusively in *cis* with M129. The wild-type *PRNP* allele in the carriers encoded V129 in 6 individuals and M129 in 4 individuals. One donor, CJD29, had a deletion of one octarepeat (1-OPRD) in the wild-type *PRNP* allele (Table 1, Figure S1).

**Table 1:**
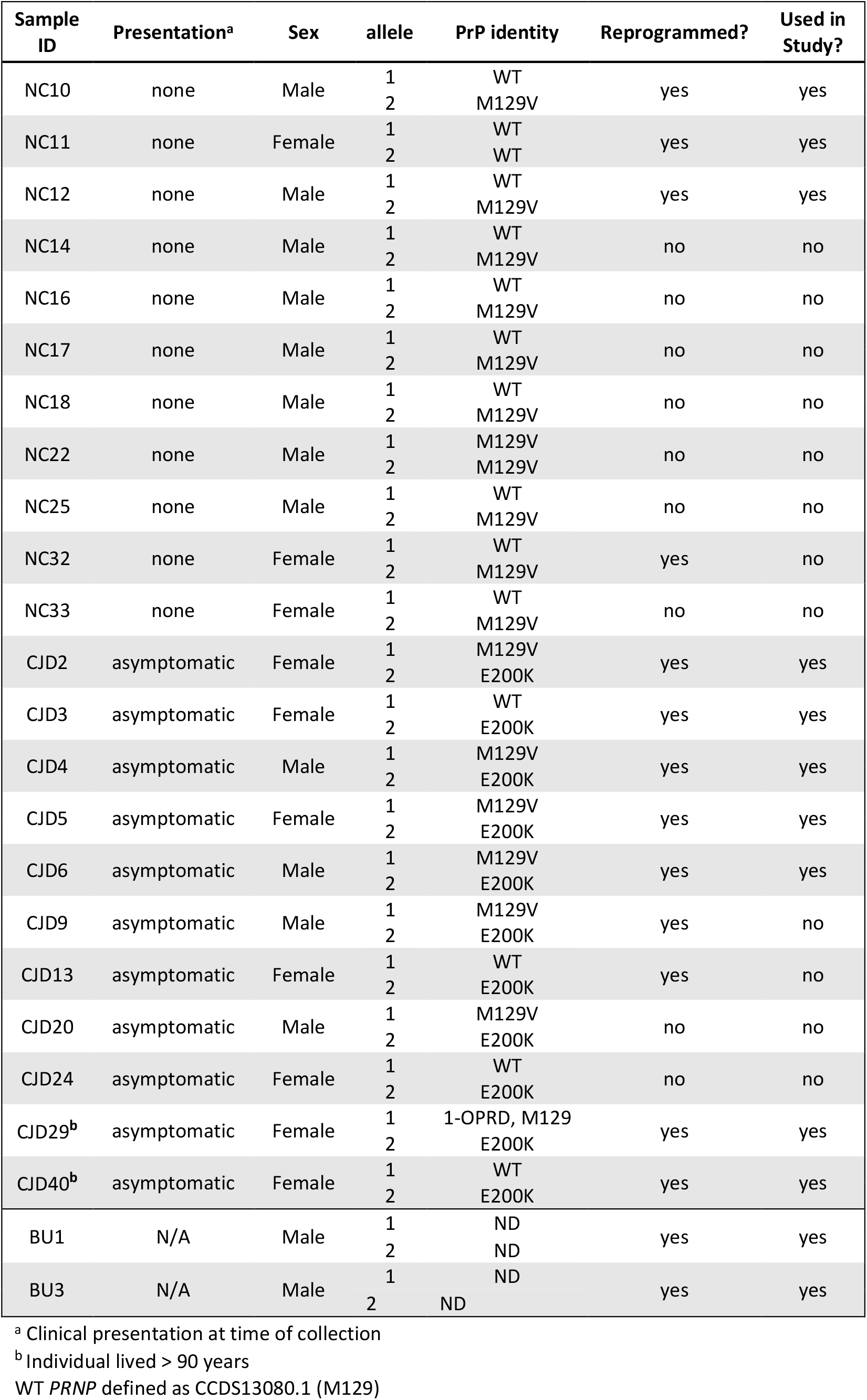
Donor characteristics.

### Whole exome sequencing

We performed whole exome sequencing of all 22 individuals. The results confirmed the three SNPs in the *PRNP* ORF observed by Sanger sequencing, including rs28933385 (c.598G>A, p.Glu200Lys); rs1799990 (c.385A>G, p.Met129Val), and rs138688873 (1-OPRD) (Figure S1). Using the whole-exome data, we interrogated eight genes in addition to *PRNP* that have been associated with disease risk or age of onset in other CJD cohorts: *CYP4X1*, a reported modifier of the age of onset of CJD-E200K (Poleggi et al., 2018); *MTMR7, STMN2, RARB*, and *NPAS2*, which have been associated with risk of variant CJD, a prion disease acquired through consumption of BSE contaminated material (Mead et al., 2009; Sanchez-Juan et al., 2012); and *GAL3ST1, STX6*, and *PDIA4*, which are associated with risk of sporadic CJD (Jones et al., 2020; Mead et al., 2019). Sequence variants in these genes could, in principle, have disease-modifying effects in our cohort, for example the absence of disease onset in CJD29 and CJD40. We found CJD associated SNPs in three of the eight genes: rs2267161 in the ORF of *GAL3ST1*; rs3747957 in the ORF of *STX6*; and rs9065 in the 3’ untranslated region of *PDIA4* (Figure S1). However, it will be necessary to obtain whole exome sequence data from a larger number of patients in this and other cohorts, particularly from those whose age of disease onset is known, to draw meaningful conclusions about genetic modifiers of E200K-associated prion disease.

### Generation of an iPSC library and differentiation into cortical neurons

Peripheral blood mononuclear cells (PBMCs) were collected and expanded *in vitro* using established procedures (Sommer et al., 2009). These cells were reprogrammed to generate iPSC lines in the absence of serum or a xenogeneic feeder layer (Figure 1A) (Park and Mostoslavsky, 2018). Expression of TRA-1-81, a marker of pluripotency, was assessed by immunofluorescence staining (Figure 1B). Next, we differentiated 12 of these iPSC lines, including two control lines derived from unrelated individuals (BU1 and BU3; Table 1), into cortical pyramidal neurons through dual SMAD inhibition (Shi et al., 2012). We adopted this differentiation protocol since it is widely used and relatively simple to perform; resulting in a relatively low percentage of astrocytes (<10%) and because the cerebral cortex is a prominent site of neuropathology in E200K CJD patients (Kovacs *et al*., 2011). iPSCs were differentiated in a stepwise manner (Figure 2A) going first through a neural stem cell stage (Figure Ci) before entering a progenitor phase within cellular aggregates known as rosettes which are positive for primary progenitor markers Pax6 and Nestin (Figure 2Cii, vi). Further neuronal induction leads to secondary progenitors, as evidenced by positive staining for TBR2 (Figure 2Cix). In the final stage of differentiation, these cells progress into mature neurons which are capable of generating Na^+^/K^+^-dependent action potentials (Figure 2B). The presence of fully differentiated neurons is further evidenced by the appearance of typical pyramidal morphology including long neuronal processes (Figure 2Civ), dendritic spine formation (Figure 2Cv), and increased expression of neuronal markers Tuj1, Map2, Vglut1, and Satb2 (Figure 2Cvii, viii, x, xi).

**Figure 1:**
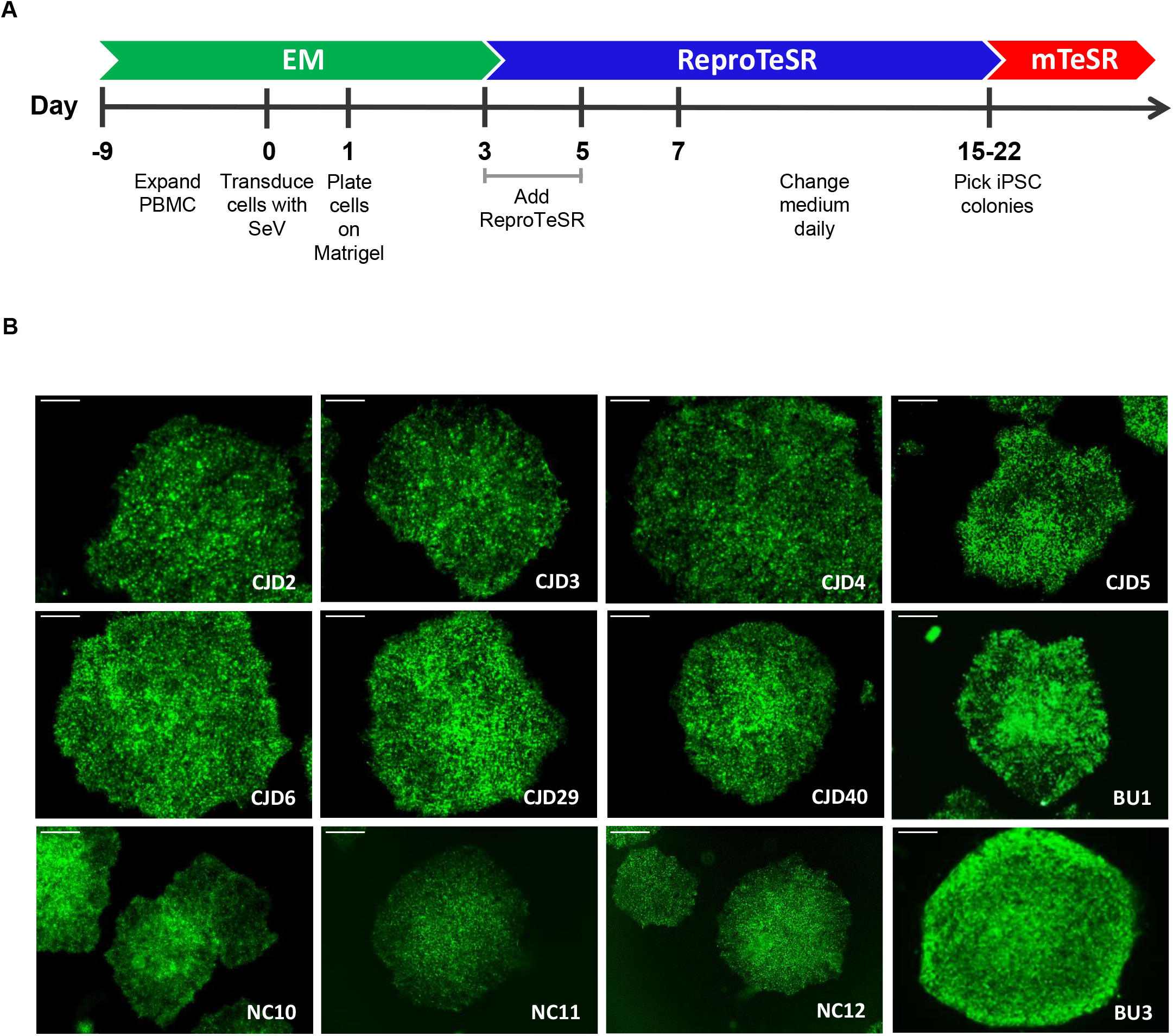
Generation of an iPSC library from patients harboring the E200K PrP mutation. **(A)** Schematic representation of the reprograming of peripheral blood mononuclear cells (PBMC) to induced pluripotent stem cells (iPSC). Sev, Sendai virus. **(B)** Representative immunofluorescence staining of TRA-1-81 from five non-carrier iPSC lines (NC10, NC11, NC12, BU1, BU3) and seven iPSC lines carrying the E200K mutation (CJD2, CJD3, CJD4, CJD5, CJD6, CJD29, CJD40) that were used in this study. Scale bars = 100 μm.

**Figure 2:**
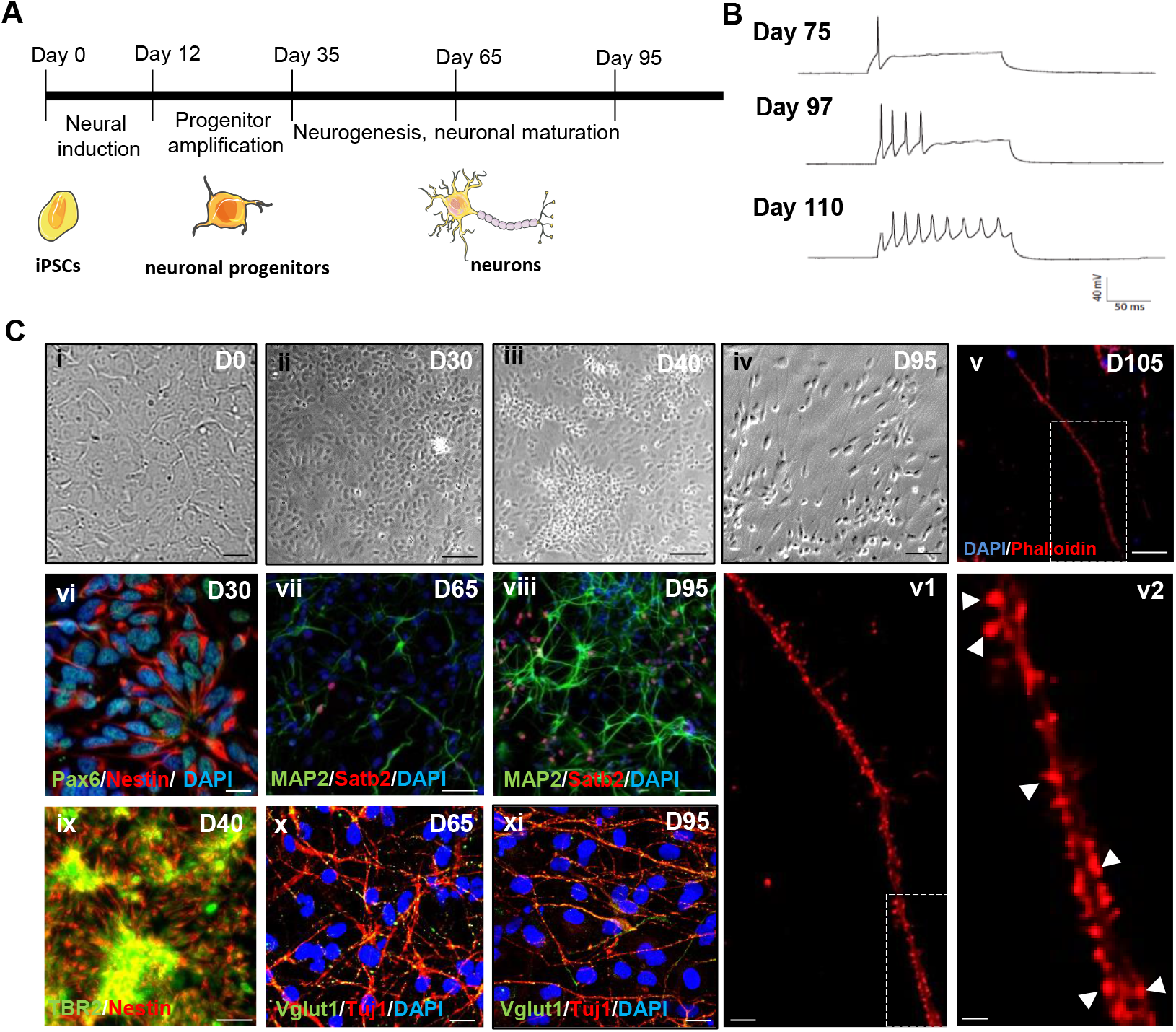
Differentiation of human iPSCs to pyramidal cortical neurons. **(A)** Schematic representation of iPSC differentiation into neuronal progenitors and neurons. **(B)** Example of electrophysiological properties of neurons at different stages of terminal differentiation. Typical single-cell, patch-clamp recordings from neurons at a range of ages following neuronal differentiation. Young neurons (day 75), fire a single action potential following current injection. As neurons mature, they progress from firing a short burst of action potentials in response to current injection (day 97) to sustained action potential firing (day 110). **(C i-v)** Morphological change in iPSCs cultures during neuronal induction. **(i)** Confluent monolayer of iPSCs ready for neural induction, **(ii)** iPSC-derived progenitor cells after neuronal induction, **(iii)** neurogenesis stage, **(iv)** neurons at day 95, **(v)** immunofluorescence staining for F-actin visualizes dendritic spine morphology of mature neurons at day 105. Each boxed region in panels **v** and **v1** is shown at higher magnification in the smaller panels to the bottom and right respectively. **(vi, ix)** Immunofluorescence staining for different progenitor makers. **(vi)** Pax6^+^/ Nestin^+^ primary progenitor cells forming a rosette structure at day 29, **(ix)** TBR2^+^/Nestin^+^ secondary progenitor cells at day 40. **(vii, viii, x, xi)** Immunofluorescence staining for different classes of excitatory neurons. The glutamatergic identity of young **(vii, x)** and mature neurons **(viii, xi)** was confirmed by immunofluorescent staining for the vesicular glutamate transporter 1 (Vglut1), neuron specific tubulin (Tuj1), dendritic marker (Map2) and cortical upper-layer neuron marker (Satb2). Nuclei in **v-xi** are stained with DAPI. These images were acquired from a differentiation of line BU1. Scale bars = 50 μm (ii, iii, iv, v, ix); 10 μm (I, vi, x, xi, v1); 1 μm (v2).

### E200K PrP from human iPSC derived neurons has some PrP^Sc^-like properties

PrP^C^ is widely expressed in the CNS and is particularly abundant in neurons, where its expression level increases during development and neuronal differentiation (Adle-Biassette et al., 2006; Steele et al., 2006; Taraboulos et al., 1992; Tremblay et al., 2007). We first examined the expression level of PrP^C^ by western blotting, revealing a similar level of PrP^C^ expression across all lines of iPSC-derived neurons at 95 days post differentiation (Figure 3A).

**Figure 3:**
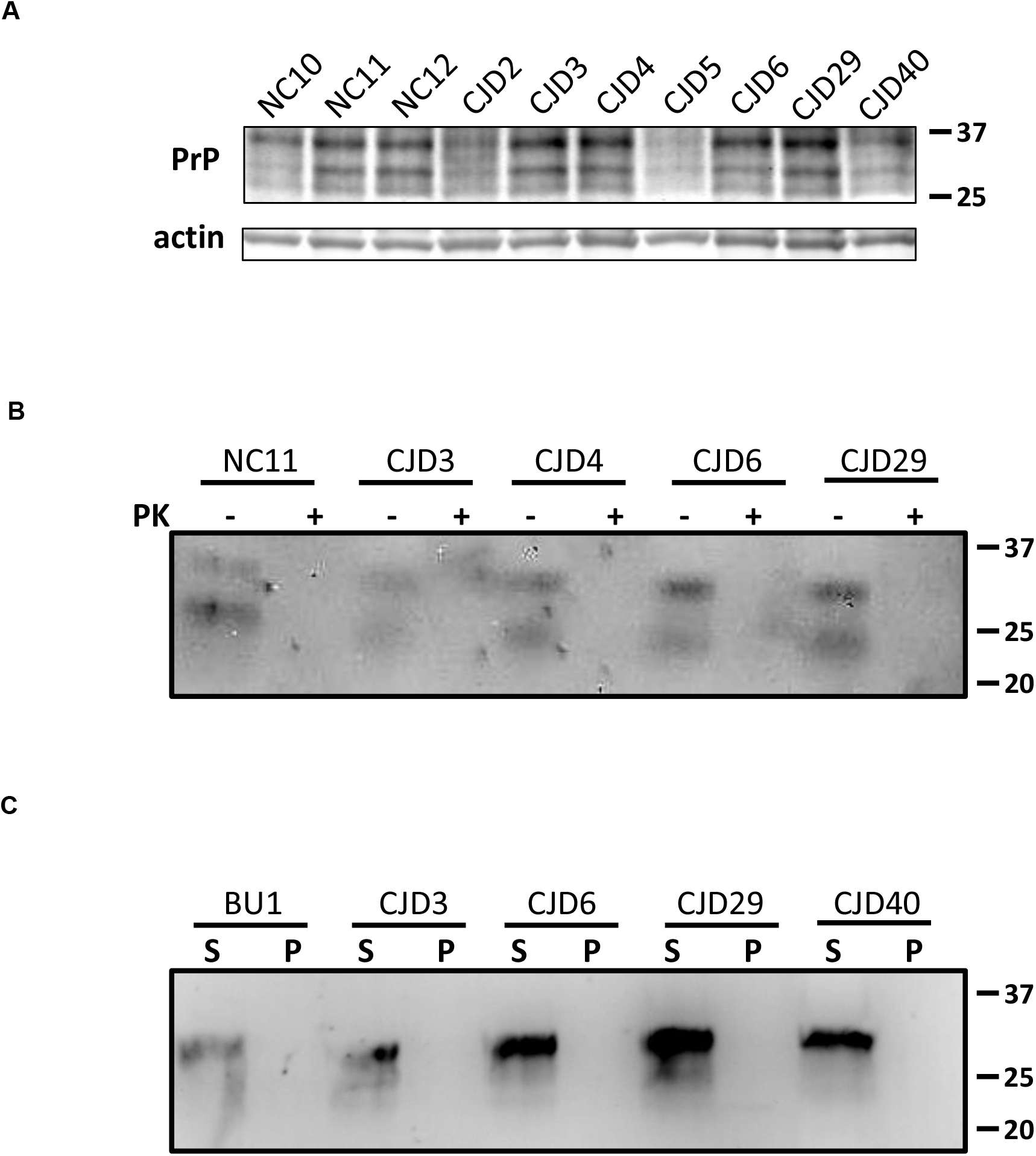
Biochemical properties of PrP expressed by iPSC derived cortical neurons. **(A)** Expression level of PrP is similar among lines of IPSCs-derived neurons. Western blotting analysis for total PrP levels in lysates from neurons at 95 days post differentiation. The anti-PrP antibody D18 was used. The three bands represent di- mono- and un-glycosylated forms of PrP, in addition to various physiological proteolytic cleavage products. **(B)** Exposure of neuronal lysates to proteinase K. 50 μg of total protein was exposed to 2.5 μg/ml PK for 1 hour at 37 °C with orbital shaking at 750 rpm. The blot was probed with the anti-PrP antibody D18. Multiple immunoreactive species are again due to differential glycosylation status of PrP. **(C)** Detergent extracts were centrifuged at 186 000 x g for 1 hour at 4 °C. Soluble and insoluble fractions were collected and analyzed by western blot. The blot was probed with the anti-PrP antibody D18.

Protease-resistant PrP^Sc^ is readily detected in the post-mortem brains of CJD-E200K patients (Kovacs *et al*., 2011), symptomatic transgenic mice expressing the equivalent mutation in the context of a mouse/human chimeric PrP (Friedman-Levi *et al*., 2011) and, to a lesser extent, in CHO cells overexpressing the mouse homologue (E199K) (Lehmann and Harris, 1996a; b). Accordingly, we performed a number of biochemical assays to detect pathologically folded PrP. First, we examined the proteinase K (PK) resistance of PrP in neuronal lysates by western blotting. Consistent with previous studies using fibroblasts derived from patient skin biopsy or cerebral organoids, we were unable to detect PK resistant PrP in neuronal lysates at 95 days post differentiation using 2.5 μg/ml of PK (Figure 3B) (Foliaki *et al*., 2020; Rosenmann et al., 2001). Next, we assessed the detergent solubility of PrP as this characteristic is known to be observable before the emergence of protease resistance (Daude et al., 1997). However, following centrifugation at 120 000 x g for 1 hour in the presence of 0.5% Triton X-100 and sodium deoxycholate, we were unable to detect PrP in the insoluble fraction (Figure 3C).

We reasoned that pathological PrP may be present in these cells, but at levels too low to be appreciable by western blotting. To investigate this possibility, we adopted an immunocytochemical assay to visualize abnormally folded PrP. The amyloid-binding dye thioflavin S (ThS) was used to visualize β-sheet-rich protein deposits, along with immunostaining to detect PrP, thus revealing pathologically folded PrP as ThS and antibody positive puncta. As a positive control, we used mouse neuroblastoma cells (N2a) that are chronically infected with Rocky Mountain Lab (RML; a mouse-adapted sheep prion strain) prions (ScN2a) (Figure 4A). Areas of co-localization between PrP and ThS were detected, which were absent in uninfected N2a cells (Figure 4A, −PK, white arrowheads). This co-localization became more pronounced upon exposure to PK to digest residual PrP^C^ (Figure 4A, +PK, white arrowhead). We found that neurons from E200K carriers contained small but detectable numbers of PrP+/ThS+ puncta after PK treatment that were not present in neurons from non-carriers (NC) or control individuals (BU) (p=0.0017 (BU + NC vs. E200K) and p=0.02 (NC vs. E200K)) (Figure 4C, D).

**Figure 4:**
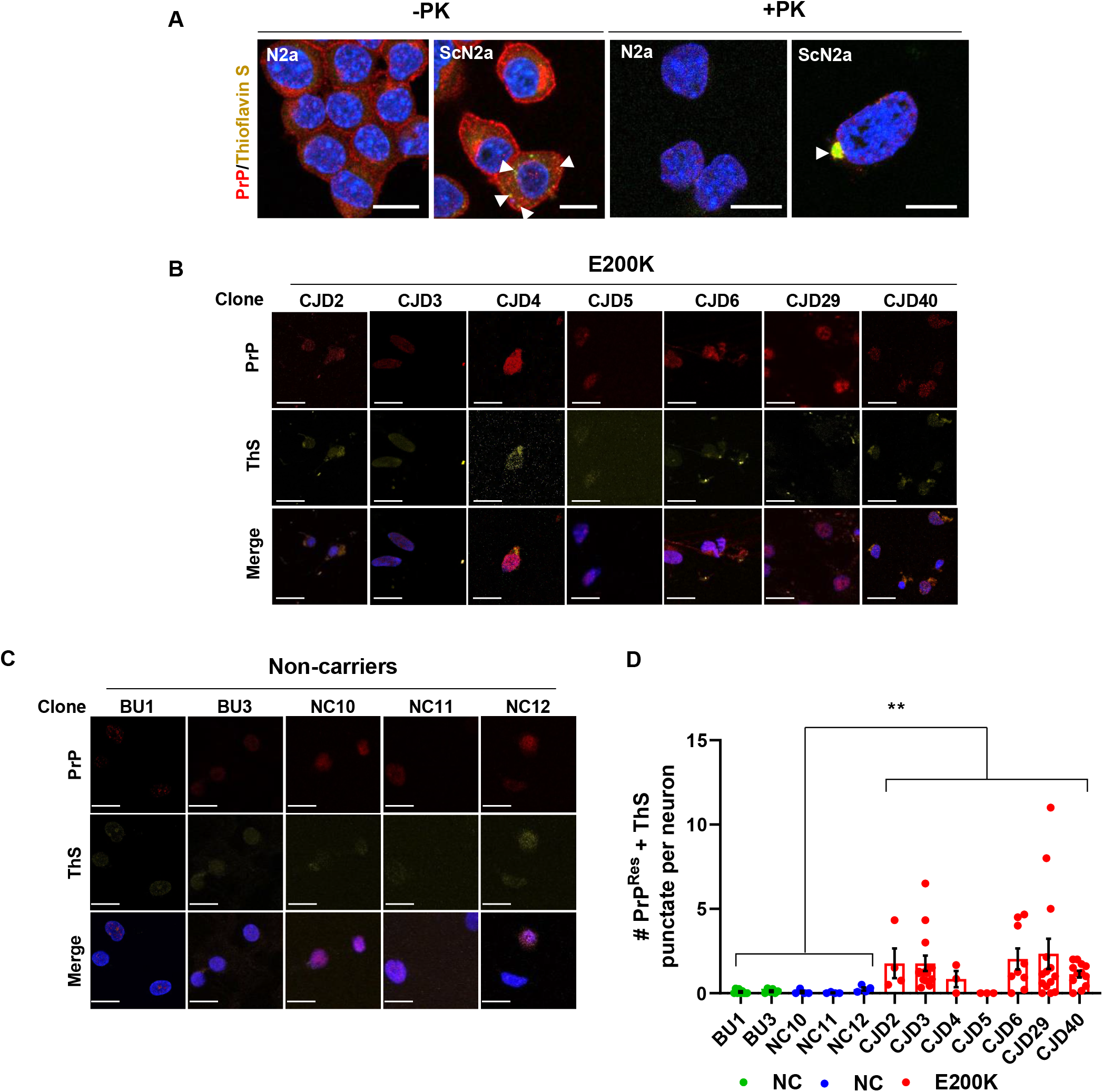
iPSC-derived neurons from E200K carriers accumulate abnormal forms of PrP. Representative images of immunofluorescent staining of PrP (D18 antibody; red) and ThS (yellow). **(A)** Uninfected N2a and ScN2a cells with and without exposure to PK. Arrows indicate accumulations of misfolded PrP. **(B)** iPSC derived neurons from individuals carrying the E200K mutation following exposure to PK. **(C)** iPSC derived neurons from non-carriers stained following exposure to PK. **(D)** Quantification of the number of PrP and ThS positive puncta following exposure to PK from B and C. Non-carriers unrelated to the kindred are indicated in green, non-carriers in the kindred in blue and E200K carriers in red. p=0.0017 (BU + NC vs. E200K) and 0.02 (NC vs. E200K). n=fields observed. BU1 n=9; BU3 n=6, NC10 n=4; NC11 n=4; NC12 n=4; CJD2 n=4; CJD3 n=14; CJD4 n=3; CJD5 n=3; CJD6 n=9; CJD29 n=14; CJD40 n=12. Mean +/- SEM is plotted. Scale bars = 10 μm.

To determine the seeding competence of misfolded PrP, we turned to Real-Time Quaking-Induced Conversion (RT-QuIC). This assay monitors PrP aggregation by measuring thioflavin T fluorescence, which increases in the presence of amyloid structures. Bank vole PrP was chosen as the recombinant substrate, as it can be seeded by a wide range of prion species and strains, and in particular, is readily able to detect CJD-E200K prions (Orrú et al., 2015). RT-QuIC is extremely sensitive to the presence of prion seeds; in our hands, we can robustly detect seeding activity in RML infected brain homogenate up to a dilution of 10^−7^ (Figure 5A) and from lysates of ScN2a cells using as little as 2 pg of total protein (Figure 5B). When used to assay 2 ng of iPSC derived neuronal cell lysate, we were unable to reliably detect seeding activity (Figure 5C). One of five replicate wells containing CJD29 lysate returned a positive result, which could indicate the presence of some seed competent PrP in this lysate near the limit of detection of the RT-QuIC assay. Interestingly, this is the line that displayed the greatest number of ThS+/PrP+ puncta (Figure 4B, D).

**Figure 5:**
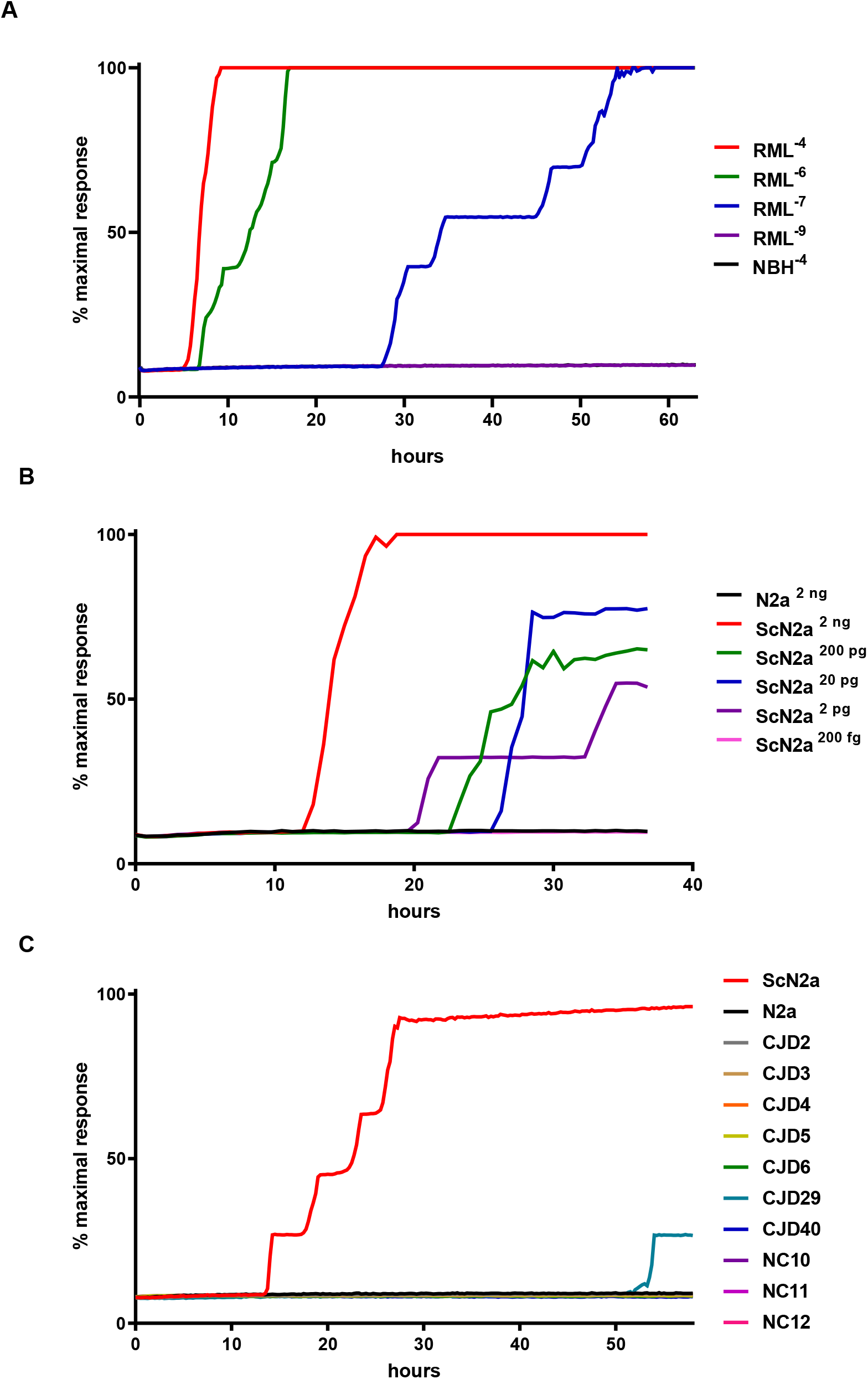
Seeding activity of cell lysates of human iPSC derived neurons. Seeding activity is assessed using the Real Time-Quaking Induced Conversion (RTQuIC) assay. **(A)** Sensitivity of the assay is demonstrated by a positive result using a 10^−7^ dilution of brain homogenate from a mouse at end-stage RML prion infection. n = 5 wells per sample. A 10^−4^ dilution of uninfected, normal brain homogenate (NBH) is used as a negative control. **(B)** RT-QuIC is capable of detecting seeding activity in as little as 2 pg of total protein from lysates of chronically RML infected N2a cells (ScN2a). Uninfected N2a cell lysate is used as a negative control. n = 5 wells per sample. **(C)** No seeding activity was detected in lysates of 95 day post differentiation iPSC derived cortical neurons. n = 5 wells seeded with 2 ng protein per sample. All experiments were performed using full length (23-230, M109) recombinant bank vole PrP (accession no. AF367624). Mean thioflavin T fluorescence as a percentage of maximum signal is plotted against time in hours.

### E200K neurons show a trend toward abnormal tau accumulation

Abnormally phosphorylated, intraneuronal deposits of tau are frequently observed by immunohistochemical staining in the brains of E200K CJD patients (Kovacs *et al*., 2011). In some cases, these phosphorylated tau species can also be observed by western blotting. We investigated whether these features could be observed in iPSC-derived neurons from E200K carriers. The levels of phospho-tau (T231) in relation to total tau by western blot was not significantly different between E200K and NC neurons (Figure S2A, B). Immunofluorescence staining revealed that E200K neurons displayed a trend toward increased numbers of p-tauT231-positive puncta compared to NC neurons, although this difference did not reach statistical significance (p = 0.07) (Figure S2C, D).

### E200K neurons show synaptic abnormalities

We next turned our attention to characterizing the synapses formed by these iPSC derived neuronal cultures. N-Methyl-D-aspartic acid receptors (NMDARs), the primary mediators of excitatory postsynaptic signaling, are stabilized at the synaptic site by the scaffolding protein post synaptic density 95 (PSD95) (Kornau et al., 1995). Of particular relevance to the current study is the finding that PrP^C^ may physically interact with some NMDAR subunits (Khosravani et al., 2008). Using immunofluorescence, we assessed the synaptic localization of NMDARs by staining for the obligatory NMDAR1 subunit and PSD95 (Figure 6A, B). Because the synaptic localization of NMDARs increases with development (Li et al., 2003), we performed these experiments with neurons at 95 days post differentiation, at which point they had matured and displayed an electrically excitable, neuronal phenotype (Figure 2B). To score colocalization, we graphed the pixel intensity from each fluorescent channel along the length of the dendrite, defining co-localization as areas of significant overlap (Figure 6C; dashed boxes). We observed significantly reduced co-localization of NMDAR1 and PSD95 staining in E200K neurons compared to neurons derived from non-carriers (Figure 6D; BU + NC vs. E200K. p=0.007; NC vs. E200K, p=0.04). This observation is not due to a change in density of either protein separately, as there was no significant difference between E200K and control neurons in the density of NMDAR1 or PSD95 dendritic clusters (Figure 6E, F).

**Figure 6:**
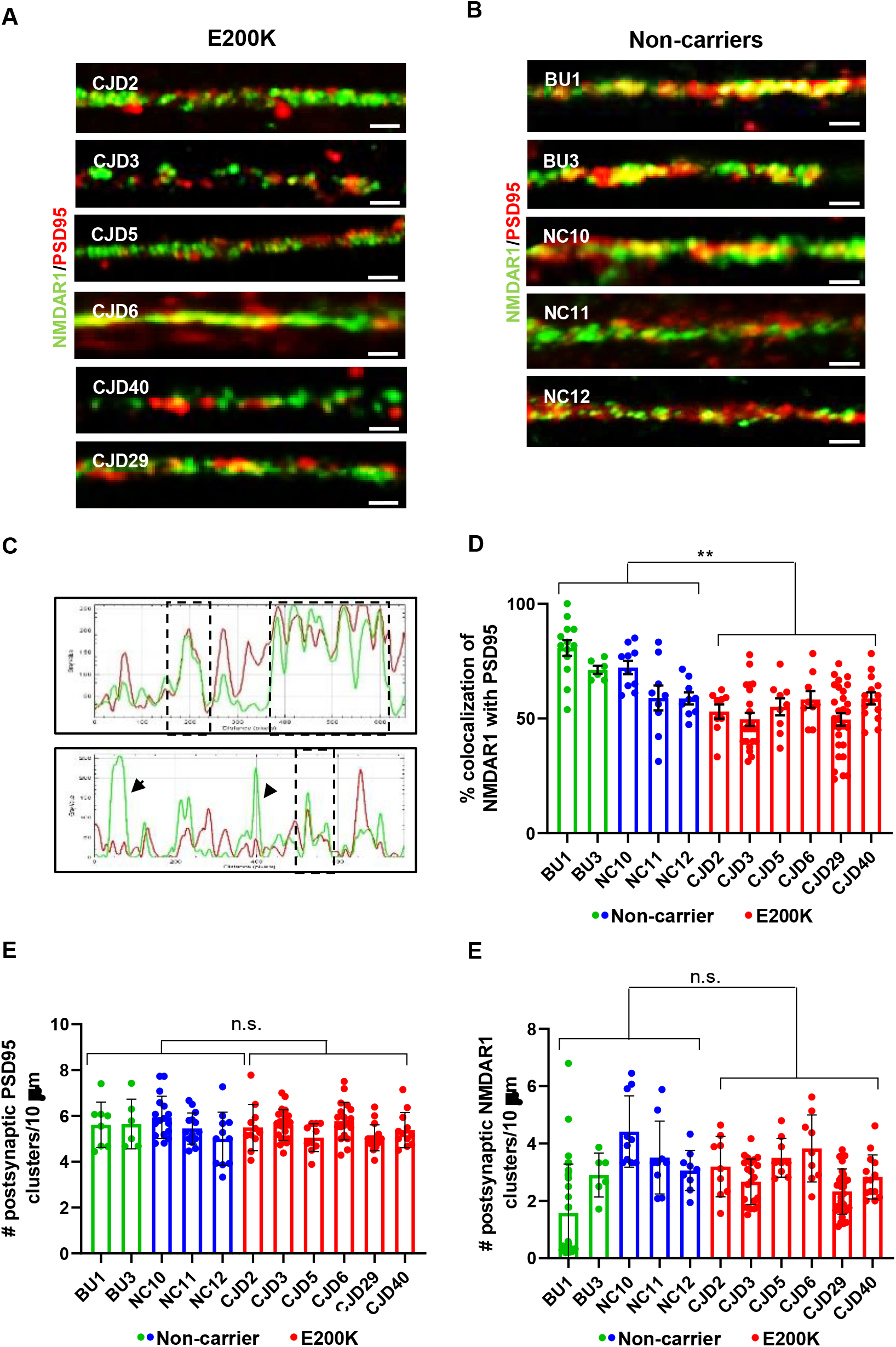
E200K PrP expressing neurons show misaligned postsynaptic markers. Representative immunofluorescence images following staining of iPSC derived cortical neurons for NMDAR1 and PSD95. NMDAR1 staining is shown in green, PSD95 staining is shown in red. **(A)** E200K carriers **(B)** non-carriers. Scale bars = 5 μm. **(C)** Graph of pixel intensities for each fluorescent channel along a representative segment of dendrite from an E200K carrier (lower panel) and a non-carrier (upper panel). Areas with significant overlap of fluorescent signal, indicated by boxes, are scored as co-localized. Black arrows indicate regions of low colocalization. **(D)** Plot showing a significant reduction in the colocalization of NMDAR1 and PSD95 along the dendritic shaft of neurons expressing E200K PrP. p=0.007 (BU + NC vs. E200K) and p=0.04 (NC vs. E200K). n=fields observed. BU1 n=13; BU3 n=6, NC10 n=10; NC11 n=9; NC12 n=9; CJD2 n=9; CJD3 n=22; CJD5 n=9; CJD6 n=9; CJD29 n=30; CJD40 n=14. **(E)** Plot of the number of postsynaptic NMDAR1 Clusters per 10 μm. Non-carriers unrelated to the kindred are indicated in green, kindred non-carriers are in blue and E200K carriers are in red. p=0.26 (BU + NC vs. E200K) and p=0.18 (NC vs. E200K). n=fields observed. BU1 n=13; BU3 n=6, NC10 n=10; NC11 n=9; NC12 n=9; CJD2 n=9; CJD3 n=22; CJD5 n=9; CJD6 n=9; CJD29 n=30; CJD40 n=14. **(F)** Plot of the number of postsynaptic PSD95 clusters per 10 μm. p=0.56 (BU + NC vs. E200K) and p=0.75 (NC vs. E200K). n=fields observed. BU1 n=8; BU3 n=6, NC10 n=17; NC11 n=14; NC12 n=11; CJD2 n=11; CJD3 n=23; CJD5 n=10; CJD6 n=19; CJD29 n=16; CJD40 n=12. Mean +/- SEM is plotted for all graphs.

## Discussion

The CJD-causing E200K mutation is the most common PrP mutation linked to genetic prion disease. In this study, we created a bank of iPSCs from a large Israeli family, which we differentiated into neurons with the characteristics of cortical projection neurons using dual SMAD signaling inhibition. In a preliminary characterization of these neurons, we detected abnormal accumulations of PrP by immunofluorescence, as well as a disturbed alignment of NMDARs and PSD95 in the postsynaptic membrane. Together, our results lay the foundation for understanding the pathogenic mechanisms underlying the E200K mutation in a cell culture system based on human neurons from patients with defined, and ongoing, clinical histories.

Given the high penetrance of the E200K mutation, and the established impact of octarepeat modifications upon disease outcomes (Chiesa et al., 1998; Lau et al., 2015), the 1-OPRD found in *trans* to E200K in the *PRNP* ORF of donor 29 is of interest for further investigation. A single octapeptide repeat deletion of *PRNP* is a relatively common polymorphism (global frequency ~0.025). It has recently been identified in an unusually late-onset case of sporadic CJD, opening the possibility that it may confer some protection (Areškevičiūtė et al., 2021; Beck et al., 2010). This is especially interesting as donor 29, along with donor 40, lived past the age of 90 years without developing CJD. As we expand this collection to include additional members of this kindred, we will continue to perform exome sequencing. Currently, all of the donors in our cohort are healthy. However, as carriers present with CJD, we will perform age of onset comparisons with these nonagenarian carriers to potentially identify disease modifying SNPs.

In previous studies, we found that PrP molecules carrying several disease-linked mutations, including E200K, exhibited biochemical properties characteristic of PrP^Sc^, including detergent-insolubility and mild protease-resistance, when expressed in transfected, immortalized cell lines in culture (Lehmann and Harris, 1996a; b). In the current study, we did not detect detergent-insoluble or protease-resistant PrP in differentiated E200K neurons using these methods. However, we did observe small numbers of PrP-containing, ThS-positive deposits by immunofluorescence in the E200K neurons that were not present in neurons derived from non-carriers. This result suggests that the mutant PrP begins a misfolding process, but does not accumulate to levels detectable by biochemical techniques during the 95 days of culture used in our study. The biochemical discrepancies between data obtained using transfected lines of immortalized cells and iPSC-derived neurons may be a reflection of the differences in expression level; iPSC-derived neurons have physiological expression levels of PrP while the transfected cell lines from previous work expressed much higher levels. Consistent with this explanation, cerebral organoids derived from E200K iPSCs also failed to accumulate detergent insoluble and protease resistant PrP (Foliaki *et al*., 2020; Smith *et al*., 2022). In contrast, insoluble, protease-resistant PrP was detected in the brains of Tg(E199K) mice, but in this case the mutant protein was also over-expressed compared to the WT protein. The low or undetectable levels of PrP^Sc^ present in our E200K neurons and in cerebral organoids correlates with the lack of seeding detected in these systems when assayed by RT-QuIC.

Accumulations of abnormally phosphorylated tau have been detected by both western blotting and immunofluorescence in the postmortem brains of CJD patients carrying the E200K mutation. While iPSC-derived E200K neurons showed a trend toward the accumulation of phospho-tau, this did not reach statistical significance; perhaps because we did not analyze a sufficient number of non-carrier neurons. Another possibility is that the tau misfolding/hyperphosphorylation process proceeds slowly when driven by physiological expression levels of mutant PrP and a longer period in culture is required for these deposits to become detectable. Increased levels of phosphorylated tau were observed in iPSC derived neurons expressing Y218N PrP after ~40 days in culture, suggesting that different mutations may have different pathological effects on neurons (Matamoros-Angles *et al*., 2018).

Glutamate receptor-dependent excitotoxicity is observed in many neurodegenerative diseases including Alzheimer’s, amyotrophic lateral sclerosis, Parkinson’s, and Huntington’s (Verma et al., 2022). Because of this, as well as our observation that exogenously applied prions cause selective, NMDAR-dependent postsynaptic degeneration in primary cultures of hippocampal neurons (Fang et al., 2018), we decided to investigate the integrity of the postsynaptic site. Interestingly, we observed a small, but statistically significant decrease in co-localization of NMDAR1 and PSD95 staining in E200K neurons, suggesting a disturbance of the postsynaptic architecture. PSD95 is a scaffolding protein that physically interacts with the cytoplasmic tails of the NR2 subunits of the NMDAR, and serves to anchor the receptors to the postsynaptic site. The lack of co-localization of PSD95 and NMDAR1 may reflect early stages of synapse disassembly. It remains to be determined whether there are electrophysiological correlates of this synaptic misalignment. Of note, PrP has been shown to interact physically with the NR2D subunit of NMDARs, raising the possibility that alterations in this function resulting from the E200K mutation could cause changes in the localization of the obligatorily associated NR1 subunit. Further experiments will be needed to delve further into the effects of E200K on synaptic structure and function.

In summary, we have generated a bank of iPSC lines from a family with an inherited prion disease due to the E200K PrP mutation. In the future, we plan to use this collection to explore key questions in the pathogenesis of inherited prion disease, including correlations of age of onset to biochemical and cellular abnormalities, the effects of the mutation on the function of iPSC-derived glial cells, the role of genetic modifiers of disease, and the efficacy of potential pharmacological interventions.

## Experimental Procedures

### Materials availability

All unique/stable reagents generated in this study are available from the lead contact with a completed Materials Transfer Agreement.

### DNA isolation and *PRNP* genotyping

gDNA was purified from PBMCs using the Qiagen DNeasy Blood and Tissue kit. The *PRNP* open reading frame was amplified by PCR using the following primers:

For: 5’-TCGGAAGCTTACGTTCTCCTCTTCATTTTGCAG-3’

Rev: 5’-CGTATCTAGAGGAAGACCTTCCTCATCCCAC-3’

PCR amplicons were digested with HindIII and XbaI and ligated into pcDNA3.1. Following transformation, clones were analyzed by Sanger sequencing. Reference sequence CCDS13080.1

### Generation of iPSCs

Derivation of iPSCs was performed as described (Park and Mostoslavsky, 2018; Somers et al., 2010; Sommer *et al*., 2009). PBMCs were expanded and reprogrammed using Sendai virus vector (CytoTune-iPS 2.0, Invitrogen) using a feeder-free system (Sommer *et al*., 2009; Staerk et al., 2010). At least 3 independent clones were established, expanded, and saved from each individual. All human subject studies were performed under signed consent and approved by the Boston University Institutional Review Board protocol H-32506.

### Differentiation of iPSC to cortical neurons

Neuronal differentiation was carried out described (Shi *et al*., 2012). Briefly, iPSC lines were cultured in mTeSR™ 1 medium (STEMCELL Technologies) on a matrigel (BD) coated dish. When the cells reached full confluence, the medium was changed to neural induction media (1:1 mixture of DMEM/F-12 GlutaMAX: Neurobasal media supplemented with 0.5x N-2 (Life Technologies), 0.5x B-27 (Life Technologies), 2.5 μg/ml insulin (Sigma), 1mM L-glutamine (Life Technologies), 0.5x NEAA (Life Technologies), 50 μM 2-mercaptoethanol (Life Technologies), 0.5 mM sodium pyruvate (Sigma), 50 U/ml penicillin and 50 mg/ml streptomycin (Life Technologies)) supplemented with 10 μM SB431542 (Tocris) and 1 μM Dorsomorphin (Tocris). After forming a dense neuro-epithelial sheet. Cells were passaged with dispase (STEMCELL Technologies) to two laminin (1-ug/ml) coated wells. Following cell reattachment, the medium was changed to neural maintenance medium. FGF2 was added the following day to a concentration of 20 ng/ml. Cells were split 1:2 with dispase when rosettes started to meet, or if neural crest cells began to appear (typically after 4 days of FGF2 treatment). Careful dispase passaging left undifferentiated cells attached, but allowed neural rosettes to lift off the substrate. On day 25 after induction (±1 day), cells were dissociated to single cells with Accutase™ (STEMCELL Technologies) at a ratio of 1:1 into new laminin coated wells. Cells were passaged for the final time around day 35 onto poly-L-ornithine/ laminin coated vessels. Media was changed daily, and mature neuronal samples are collected at day 95 of differentiation for biochemical analysis.

### Electrophysiology

Recordings were made from iPSC-derived neurons 60-110 days after induction. Whole-cell patch clamp recordings were collected using standard techniques. Pipettes were pulled from borosilicate glass and polished to an open resistance of 2–5 megaohms. Experiments were conducted at room temperature with the following solutions: internal, 140 mM Cs-gluconate, 5 mM CsCl, 4 mM MgATP, 1 mM Na_2_GTP, 10 mM EGTA, and 10 mM HEPES (pH 7.4 with CsOH); external, 150 mM NaCl, 4 mM KCl, 2 mM CaCl_2_, 2 mM MgCl_2_, 10 mM glucose, and 10 mM HEPES (pH 7.4 with NaOH). Signals were collected from a Multiclamp 700B amplifier (Molecular Devices), digitized with a Digidata 1550A interface (Axon Instruments), and saved to disc for analysis with PClamp 10 software.

### Immunocytochemical analysis

iPSCs and differentiated neurons were grown on coverslips coated with poly-L-ornithine/ laminin. After washing in PBS, cells were fixed using 4% (w/v) PFA (Sigma) in PBS for 15 min and permeabilized with 0.1% Triton X-100 (Sigma) in PBS for 5 min, washed and blocked in 1% BSA (Sigma) in PBS for 30 minutes. Cells were stained using the primary and secondary antibodies listed (Table S1). Neurons were mounted in Vectashield Mounting Medium and images were acquired using a Zeiss LSM 700 or 880 Laser Scanning Confocal Microscope with Airyscan using a 63x objective. Quantification was carried out on 30-50 neurons for each line.

### Western blotting

Cells were lysed with RIPA buffer (50 mM Tris base, 150 mM NaCl, 0.1% SDS, 0.5% sodium deoxycholate, 0.5% NP40) containing protease inhibitors (ThermoFisher) and a BCA assay was performed to determine protein concentration (Pierce). Samples were electrophoresed with 12% Criterion TGX precast gels (BioRad) and transferred to PVDF membranes (Millipore). Membranes were blocked with 5% milk in TBS with 0.1% tween 20 (Fisher). Membranes were incubated overnight in the indicated primary antibody and after washing were incubated in the appropriate HRP-conjugated secondary antibody, exposed to ECL (Millipore) and visualized with a Gel Doc (BioRad). Quantification was performed using ImageJ. p values were determined using the student’s t-test.

### Detergent insolubility assay

One-tenth volume of 10x detergent was added to cell lysate to a final concentration of 0.5% Triton X-100 and 0.5% sodium deoxycholate. Samples were incubated for 10 min on ice, and debris removed by spinning at 1000 g for 5 min at 4 °C. Samples were then centrifuged at 120000 x g for 40 min at 4 °C using a TLA-55 rotor. Supernatant (S) and pellet (P) samples were then analyzed by western blotting.

### Detection of pathological forms of prion protein by immunofluorescence and Thioflavin S staining

PrP^Sc^ was revealed by exposure to PK and treatment with guanidine hydrochloride to remove PrP^C^ and expose PrP^Sc^ as described (Goold et al., 2011; Veith et al., 2009). N2a and ScN2a cells were seeded to semi-confluence on glass coverslips (pretreated with

poly-L-lysine (0.2mg/ml) for 1h at RT) for 24h. iPSC derived neurons were cultured on poly-L-ornithine/ laminin coated coverslips. Cells were washed with PBS, and simultaneously fixed with freshly prepared paraformaldehyde (4% for non-PK treatment, 8% for PK treatment) and permeabilized with 0.1% Triton-X 100 for 1h at RT. Cells were then exposed to PK (5 μg/ml) for 10 min at 37 °C (stopped with 2mM PMSF (Sigma) for 15 min), as indicated. Cells were then incubated with 6M guanidine hydrochloride for 10 minutes. After blocking with 1% BSA for 30 min at RT, the cells were incubated with 0.025% of thioflavin S for 8 min and washed three times with 80% ethanol for 5 min. PrP was detected using antibody D18 in 1% BSA in PBS overnight at 4 °C followed by goat anti human-Alexa fluor 594 in 1% BSA in PBS for 45 min. Finally, cells were washed with PBS, counterstained with DAPI to reveal nuclei, and mounted in Vectashield Mounting Medium. The fluorescence was measured in at least 10 randomly chosen observation fields for each experimental condition using Zeiss LSM 880 Laser Scanning Confocal Microscope with Airyscan. Total Cell Fluorescence (CTCF) was calculated using the formula: CTCF = Integrated density – (Area of selected cell X Mean fluorescence of Background readings) using the ImageJ 2.0.0 Software (NIH). Quantification experiments were carried out independently at least three times; more than 30 neurons were counted for each condition.

### Real Time-Quaking Induced Conversion (RT-QuIC) Assay

Recombinant bank vole PrP (residues 23 to 230; Methionine at position 109; accession no. AF367624) was expressed and purified as described previously (Orrú *et al*., 2015). The concentration of recPrP was determined using a molar extinction coefficient at 280 nm of 62,005/M/cm. The protein was aliquoted and stored at −80 °C until use. RT-QuIC reactions were performed in black clear bottom 96-well plates using a reaction mixture composed of 10 mM phosphate buffer (pH 7.4), 300mM NaCl, 0.001% SDS, 1 mM EDTA, 10 μM ThT and 0.1 mg/ml PrP in a final volume of 98 μL and seeded with 2 μL of sample. Plates were sealed and incubated in a BMG Polarstar plate reader at 42 °C with cycles of 1 min of shaking (700 rpm double orbital) and 1 min of rest. ThT fluorescence measurements were taken every 15 min (450 ± 10 nm excitation and 480 ± 10 nm emission; bottom read; 20 flashes per well; manual gain of 2000; 20 s integration time). Data were analyzed using GraphPad Prism 8.

### Statistical Analysis

We statistically analyzed variables of interest by study groups (NC vs. E200K) in mixed linear models that accounted for within-line clustering resulting from the analysis of multiple observations per cell line and assume a Gaussian distribution for these dependent variables. We employed a working correlation structure of compound symmetry to address this clustering which, if left unaccounted, would have produced estimates of standard error that would have been too small, leading to the possibility of unjustifiably declaring statistical significance for differences in mean values between the study groups. We present the estimated means and standard errors from these mixed linear models. We define statistical significance where analyses have p values less than < 0.05.

## Acknowledgements

The authors would like to acknowledge the contribution of the family members, without which this work would not be possible. We would also like to acknowledge Professor Howard Cabral for assistance with statistical analysis and Dr. Byron Caughey for providing the bank vole PrP expression plasmid. Nhat T.T. Le was supported by a Warren Alpert Distinguished Scholar Award (Ltr dtd 3/28/2019), and Robert C.C. Mercer by a grant from the Department of Defense (PR201695). Work in the Mostoslavsky lab is supported by NIH Grants N0175N92020C00005 and 1R01DA051889-01 as well as a grant from the Creutzfeldt-Jakob disease Foundation. Work in the Harris lab is supported by NIH R01 NS065244. This work was also supported by NIH grant 1R21NS111499-01 to GM and DAH.

## Author Contributions

Conceptualization, Project administration, Funding acquisition and Supervision: GM & DAH; Resources: AA, GM & DAH; Data curation, Formal analysis: NTTL, RCCM, ADG, BW, PSB, RGC; Visualization: NTTL, RCCM, ADG; Investigation, Methodology: NTTL, RCCM, ADG, SP, BW, PSB, RGC; Writing – original draft: RCCM, DAH; Writing – review & editing: NTTL, RCCM, ADG, GM & DAH.

