## Supplemental Information for "NMDA receptor misalignment in iPSC-derived neurons from a multi-generational family with inherited Creutzfeldt-Jakob disease"

**Supplemental Information**
**Supplemental Figures**

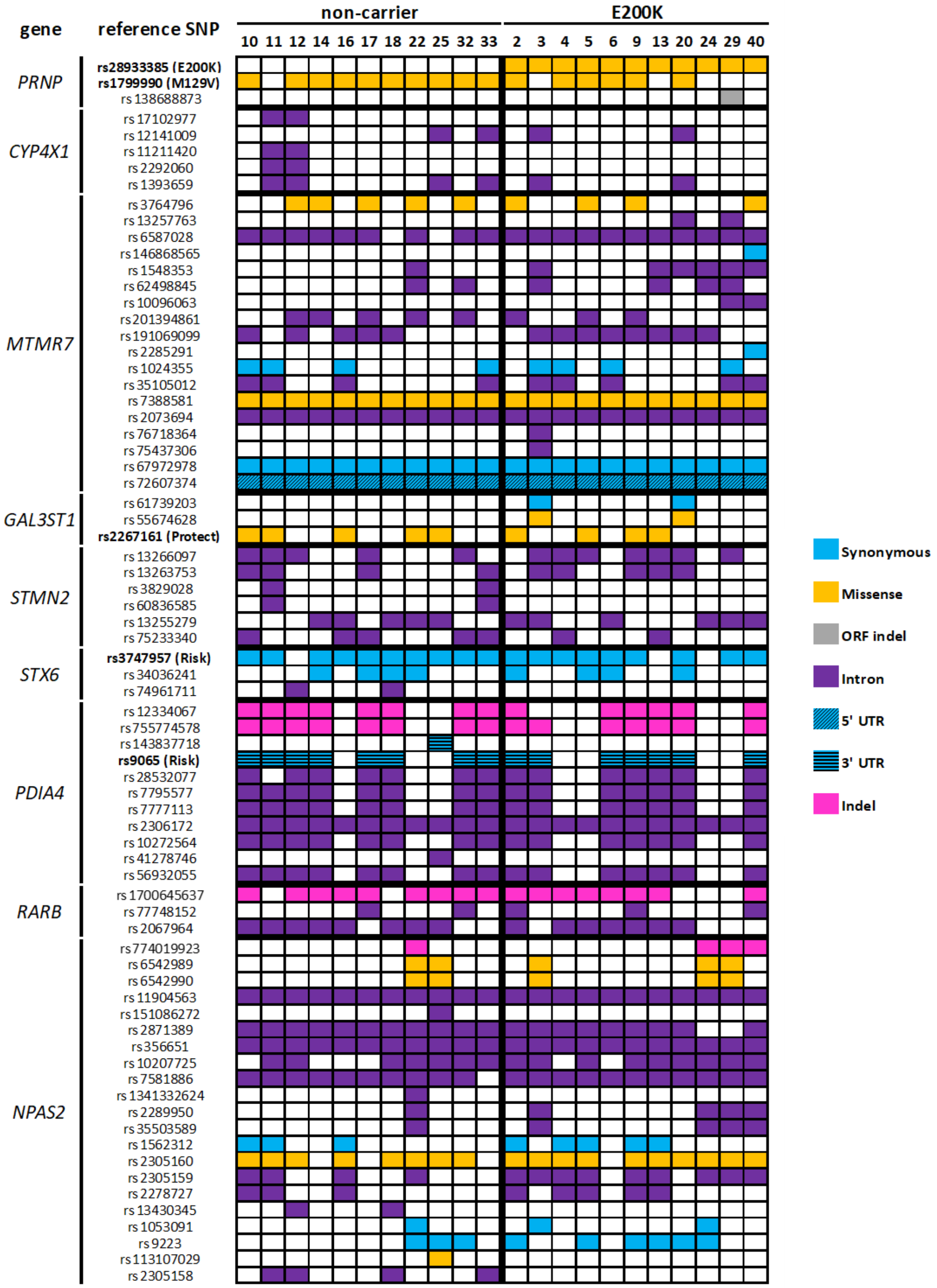


**Figure S1: Genetic variation in CJD risk loci.**
Graphical representation of exome sequencing data at nine genomic loci associated with CJD risk. The presence of single nucleotide polymorphisms (SNPs) are represented by filled in boxes. Light blue = synonymous mutation within an exon, orange = missense mutation within an exon, grey = insertion/deletion (indel) within an exon; purple = mutation within an intron, light blue with diagonal lines = 5' untranslated region mutation (5' UTR), light blue with horizontal lines = 3' untranslated region mutation (3' UTR), pink = insertion/deletion (indel) within an intron. SNPs identified in genome wide association studies of CJD are listed in bold.

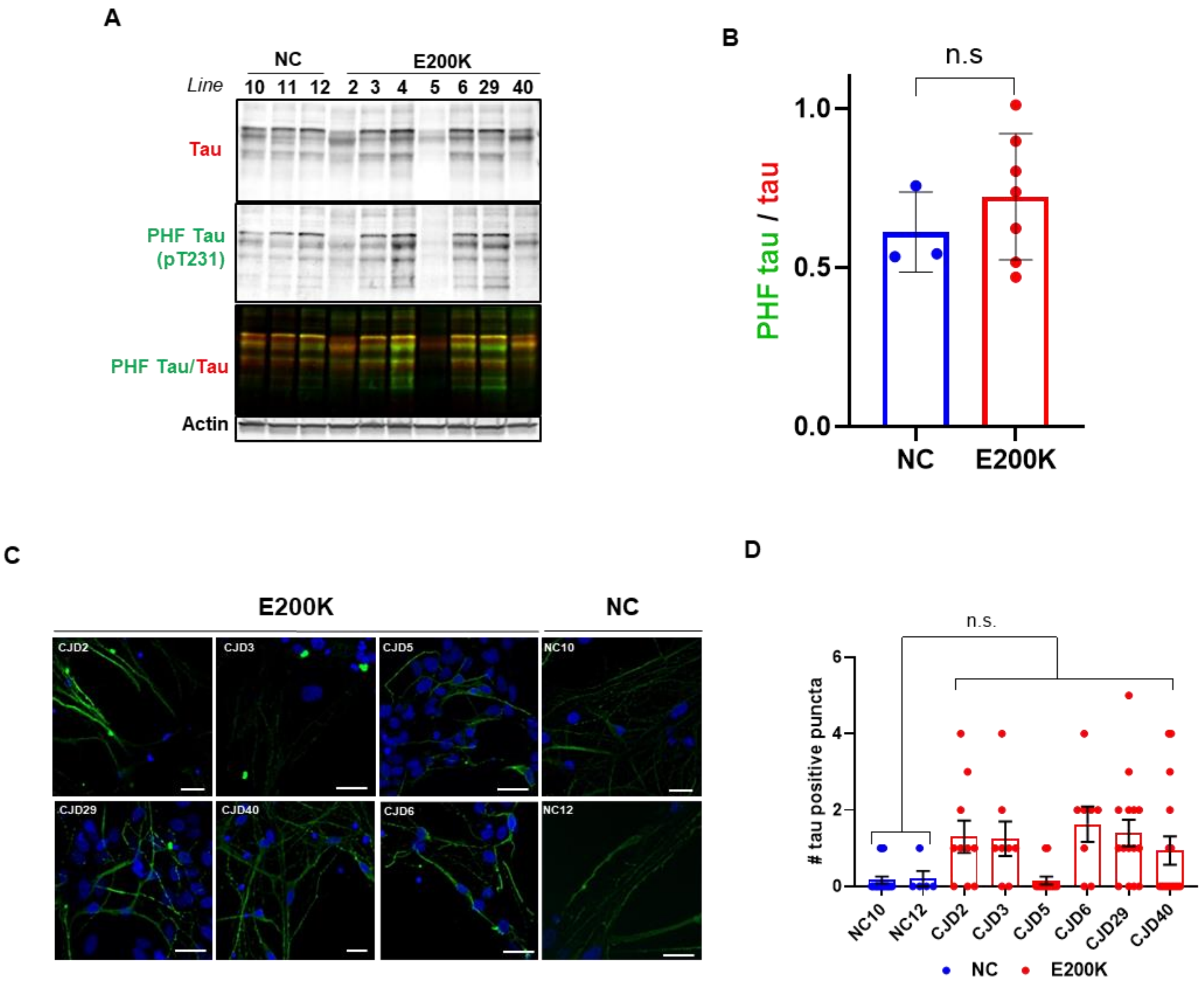


**Figure S2: Pathological tau accumulation in iPSC derived neurons**

**(A)** Western blot of neuronal lysates for total tau and PHF tau (p-tauT231). **(B)** Quantification of blots in A. One tailed students t-test comparing E200K expressing neurons (n=7) and WT control neurons (n=3) showed no significant difference, p=0.2022 **(C)** Representative images of immunofluorescence staining of tau (PHF-tau, p-tauT231) in iPSC derived cortical neurons. PHF tau staining is shown in green and DAPI in blue. **(D)** Plot of the number of PrP and tau positive puncta found in neurons derived from non-carriers and E200K carriers. p=0.07 (NC vs. E200K). n =fields observed. NC10 n=18; NC12 n=5; CJD2 n=10; CJD3 n=8; CJD5 n=13; CJD6 n=8; CJD29 n=15; CJD40 n=16. Mean +/- SEM is plotted. Scale bars = 15 µm.

**Table S1: Antibodies used in study**

| Antibody | Species | Dilution | | Company (catalogue #) |
|---|---|---|---|---|
| | | IF | WB | |
| Pax6 | Rabbit | 1:200 | - | StemCell (60094) |
| Nestin | Mouse | 1:500 | - | StemCell (60091) |
| Tbr2 | Rabbit | 1:200 | - | Abcam (ab23345) |
| Satb2 | Mouse | 1:100 | - | Abcam (ab51502) |
| Map2 | Rabbit | 1:200 | - | Abcam (ab32454) |
| Vglut1 | Rabbit | 1:300 | - | Synaptic systems (135303) |
| Tuj1 (TUBB3) | Mouse | 1:300 | - | BioLegend (801202) |
| D18 | Human | - | 1:10000 | Produced in Harris Lab |
| PHF-Tau (AT180) | Mouse | 1:100 | 1:1000 | ThermoFisher (MN1040) |
| Tau | Rabbit | 1:100 | 1:1000 | Dako (A0024) |
| NMDAR1 | Mouse | 1:100 | - | EMD Millipore (MAB363) |
| PSD95 | Rabbit | 1:100 | - | Abcam (ab18258) |
| Actin | Mouse | - | 1:5000 | Sigma-Aldrich (MAB1501) |
| Rhodamine-Phalloidin | - | 1:200 | - | ThermoFisher (R415) |
| Alexa Fluor 488 anti-mouse | - | 1:200 | - | Invitrogen (A32766) |
| Alexa Fluor 555 anti-rabbit | - | 1:200 | - | Invitrogen (A31572) |
| Alexa Fluor 594 anti-rabbit | - | 1:200 | - | Invitrogen (A32754) |
| Alexa Fluor 643 anti-mouse | - | 1:200 | - | Invitrogen (A31571) |
| Alexa Fluor 555 anti-human | - | 1:200 | - | Invitrogen (A21433) |
| IRDye 680DR anti-rabbit | - | - | 1:10000 | LI-COR (926-68073) |
| IRDye 800CW anti-mouse | - | - | 1:10000 | LI-COR (926-32212) |
| anti-goat HRP | - | - | 1:10000 | BioRad (1705047) |
| anti-rabbit HRP | - | - | 1:10000 | BioRad (1706515) |

**Supplemental Methods**

**Whole Exome Sequencing and SNP analysis**

DNA libraries were created by enzymatically shearing DNA to a mean fragment size of 200 base pairs, and a common Y-shaped adapter was ligated to all DNA libraries. Unique, asymmetric 10-base-pair barcodes were added to the DNA fragment during library amplification to facilitate multiplexed exome capture and sequencing. Equal amounts of sample were pooled before overnight exome capture, with a slightly modified version of IDT's xGen probe library. The captured DNA was PCR-amplified and quantified by quantitative PCR. The multiplexed samples were pooled and then sequenced using 75-base-pair paired-end reads with two 10-base-pair index reads on the Illumina NovaSeq 6000 platform.

Sample read mapping and variant calling, aggregation and quality control were performed using the SPB protocol. In brief, for each sample, NovaSeq WES reads are mapped with BWA MEM to the hg38 reference genome. Small variants are identified with WeCall and reported as per-sample gVCFs. These gVCFs are aggregated with GLnexus into a joint-genotyped, multi-sample project-level VCF (pVCF). SNV genotypes with read depth (DP) less than 7 and indel genotypes with read depth less than 10 are changed to no-call genotypes. After the application of the DP genotype filter, a variant-level allele-balance filter is applied, retaining only variants that meet either of the following criteria: (i) at least one homozygous variant carrier; or (ii) at least one heterozygous variant carrier with an allele balance (AB) greater than the cut-off ($AB \geq 0.15$ for SNVs and $AB \geq 0.20$ for indels). Samples showing disagreement between genetically determined and reported sex, high rates of heterozygosity or contamination (estimated with the VerifyBamId tool, specifically with a FREEMIX score > 5%), low sequence coverage (less than 80% of targeted bases achieving 20× coverage) or genetically identified sample duplicates, and WES variants discordant with genotyping chip, were excluded. *PRNP*, *RARB*, *STMN2*, *MTMR7*, *NPAS2*, *STX6*, *GAL3ST1*, *PDIA4* and *CYP4X1* gene coordinates were downloaded from ensembl. The downloaded region of gene each was extended 2kb upstream to include the promoter region of that gene. Using BEDtools (Quinlan and Hall, 2010) intersect command variants overlapping with selected genes and its promoter region were filtered from the VCF files corresponding to each of the donors.
